# Uncovering historical small mammal biodiversity among the Madrean Sky Islands

**DOI:** 10.1101/2025.09.04.674278

**Authors:** Damien C. Rivera, Nathan S. Upham

## Abstract

The Madrean Sky Island Archipelago is a system of 54 mountains with isolated woodland habitat above 1,600 meters, primarily in the Sonoran Desert. These mountains harbor a wide variety of native small mammals spanning 11 families of bats, rodents, and shrews. Improved understanding of Madrean Sky Island biodiversity should advance studies of biogeography, phylogeny, host-symbiont interactions, and community assembly in this ecoregion. However, which species are found in each sky island and how their populations are genetically related remain open questions. To establish the current knowledge baseline, we used georeferenced voucher specimens to summarize the extent and timing of past collecting efforts for small mammals in woodland habitats across the Madrean Sky Islands. In total, at least 88 species (35 bats, 50 rodents, 3 shrews) from 9,540 specimens were collected from 1884–2023. Of these specimens, 79% come from 5 sky islands (Chiricahuas, Pinaleños, Huachucas, Animas, and Santa Catalinas) and at least 23 sky islands in the Madrean system have no recorded specimens. Mexico’s 25 sky islands are mostly unsampled (only the San Luis, Sierra dos Ajos, and Sierra La Mariquita have any georeferenced specimens) and several of Arizona’s larger sky islands have fewer than 40 specimens (Galiuros, Canelo Hills, Santa Teresas, Mules, and Dragoons). Most small mammal specimens (88%) were collected in 1980 or earlier without prioritizing frozen tissues for DNA/RNA preservation. Including 3,946 non-georeferenced specimens may add sampling for another 6 sky islands and 6 species, but georeferencing confirmation is needed and major patterns are unchanged. This distributional summary is the current basis for all derived biodiversity knowledge of Madrean Sky Island small mammals, illustrating clear gaps regarding most species of woodland-dwelling bats, rodents, and shrews. There is a critical need for future fieldwork and voucher specimen preservation (including of flash-frozen tissues) in this region, both to monitor changes compared to historical sampling and to test biodiversity assumptions on unsampled mountains.

## Introduction

> “Mountains—those islands in the sky surrounded by a sea of desert.”

> – Edward Abbey (*Desert Solitaire,* 1968: 129)

Envisioning forested desert mountains as ‘sky islands’ dates back to Heald’s (1967) account of Arizona’s Chiricahua Mountains, and Abbey’s (1968) ode to Utah’s La Sal Mountains. The metaphor has inspired generations of naturalists, contributing to the foundations of metapopulation ecology and habitat island biogeography (reviewed in Matthews 2021). The Madrean Sky Island Archipelago, containing the Chiricahuas among other mountains, is globally one of the most extensive of the so-called ‘stepping stone’ archipelagos that connect two mountain chains (Warshall 1995). Spanning between the Colorado Plateau and the Sierra Madre Occidental, the Madrean Archipelago contains somewhere between 40 (Warshall 1995) and 65 (Moore et al. 2013) forested mountains, each one isolated from others by the surrounding Sonoran and Chihuahuan Deserts. Here we present a 54-mountain definition of the Madrean Sky Islands designed for studying woodland-dwelling species relative to the surrounding ‘sea’ of greater aridity (desert grassland and desert scrub; Fig. 1, Table 1).

**Fig 1.**
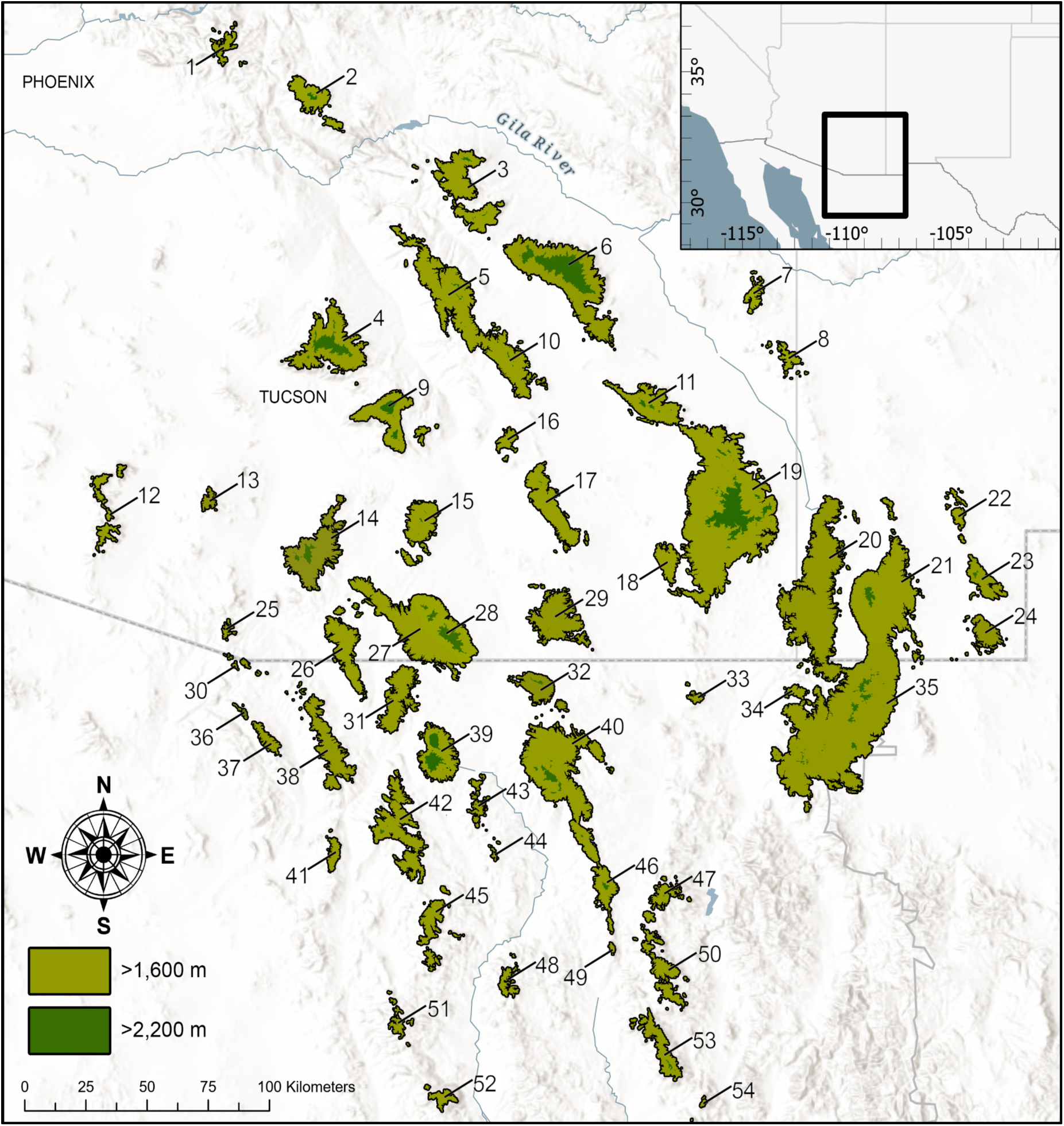
Map of the Madrean Sky Island region showing the 54 mountains defined in this study, with numbered labels corresponding to Table 1. The Colorado Plateau lies directly North of the figure and the Sierra Madre Occidental lies South/Southeast of the figure.

**Table 1.** List of all 54 mountains in the Madrean Sky Islands region as defined in this study, with the label numbers corresponding to Figure 1.

| Label | Mountain | Area above 1,600<br>meters (km<sup>2</sup>) | Maximum elevation<br>(m) |
| --- | --- | --- | --- |
| 1 | Superstition | 43.28 | 1902 |
| 2 | Pinal | 144.36 | 2392 |
| 3 | Santa Teresa | 327.76 | 2497 |
| 4 | Santa Catalina | 414.78 | 2806 |
| 5 | Galiuro | 501.57 | 2322 |
| 6 | Pinaleño | 747.2 | 3267 |
| 7 | Peloncillo North | 40.3 | 1982 |
| 8 | Peloncillo Central | 38.11 | 1982 |
| 9 | Rincon | 264.1 | 2646 |
| 10 | Winchester | 234.28 | 2315 |
| 11 | Dos Cabezas | 218.99 | 2485 |
| 12 | Baboquivari | 69.59 | 2262 |
| 13 | Sierrita | 24.12 | 1887 |
| 14 | Santa Rita | 341.77 | 2869 |
| 15 | Whetstone | 183.1 | 2331 |
| 16 | Little Dragoon | 39.81 | 2037 |
| 17 | Dragoon | 252.4 | 2275 |
| 18 | Swisshelm | 83.45 | 2181 |
| 19 | Chiricahua | 1769.87 | 2990 |
| 20 | Peloncillo South | 903.55 | 2095 |
| 21 | Animas | 739.77 | 2598 |
| 22 | Little Hatchet | 35.47 | 2017 |
| 23 | Big Hatchet | 109.32 | 2542 |
| 24 | Alamo Hueco | 89.17 | 2090 |
| 25 | Atascosa | 11.09 | 1948 |
| 26 | Patagonia | 201.43 | 2200 |

| **Label** | **Mountain** | **Area above 1,600**<br>**meters (km<sup>2</sup>)** | **Maximum elevation**<br>**(m)** |
| --- | --- | --- | --- |
| 27 | Canelo Hills | 416.09 | 1968 |
| 28 | Huachuca | 357.84 | 2875 |
| 29 | Mule | 265.56 | 2241 |
| 30 | Pajarito | 8.89 | 1800 |
| 31 | Sierra Chivato | 192.43 | 2195 |
| 32 | Sierra San Jose | 123.24 | 2539 |
| 33 | Sierra La Ceniza | 13.61 | 1822 |
| 34 | Sierra de Embudos | 24.31 | 1921 |
| 35 | San Luis | 1722.44 | 2515 |
| 36 | Sierra Avispas | 5.39 | 1746 |
| 37 | Sierra Cibuta | 43.72 | 2079 |
| 38 | Sierra El Pinito West | 52.24 | 2264 |
| 39 | Sierra La Mariquita | 239.5 | 2500 |
| 40 | Sierra de los Ajos | 719.64 | 2616 |
| 41 | Sierra La Madera | 38.22 | 2044 |
| 42 | Sierra Azul | 296.54 | 2452 |
| 43 | Sierra El Manzanal | 28.7 | 1919 |
| 44 | Cerro Bacoachi | 3.79 | 1707 |
| 45 | Sierra San Antonio | 112.39 | 2213 |
| 46 | Sierra Buenos Aires | 94.7 | 2474 |
| 47 | Sierra El Pinito East | 37.42 | 2230 |
| 48 | Cerro El Bellotal | 35.63 | 1942 |
| 49 | Sierra Purica | 11.5 | 1925 |
| 50 | Sierra La Sandia | 137.04 | 2179 |
| 51 | Sierra El Jacaral | 224.8 | 1908 |
| 52 | Sierra Aconchi | 37.43 | 2190 |
| 53 | Sierra de Oposura | 133.79 | 2345 |
| 54 | Sierra de Las Guijas | 4.11 | 1823 |

Intersecting biomes make the Madrean Sky Islands a biodiversity hotspot for plants, insects, birds, mammals, and other taxa with biogeographic origins in both Nearctic and Neotropical realms (Spector 2002; Deyo et al. 2013). They have been hailed as a ‘natural laboratory’ for testing processes of community assembly and demographic change, with replicated elevational gradients in a historically dynamic environment of glacial-interglacial cycles (Van Devender 1990, Warshall 1995, Moore et al. 2013). However, present-day distributional knowledge of the Madrean biota remains highly incomplete. Efforts to characterize species distributions have focused on flora or fauna of the more accessible mountains (e.g., Santa Catalinas—Lange 1960; Moore et al. 2013; Chiricahuas—Cahalane 1939; Duncan 1990; Huachucas—Hoffmeister and Goodpasters 1954; Bowers and McLaughlin 1996). A few individual species have also been studied across multiple mountains (e.g., Mexican Woodrat, *Neotoma mexicana*—Sullivan 1994; Canyon Tree Frog, *Hyla arenicolor*—Barber 1999; Mexican Jay, *Aphelocoma wollweberi—* McCormack et al. 2008; Yarrow’s Spiny Lizard, *Sceloperous jarrovii*—Wiens et al. 2019; American Black Bear, *Ursus americanus*—Atwood et al. 2011; Gould et al. 2022). Given recent losses of forested habitats in the Madrean Archipelago from fire, logging, and climate change (Williams et al. 2010; Yanahan and Moore 2019), there is growing urgency for a more integrative multi-species and multi-mountain understanding of the region’s biodiversity. Identifying gaps in the historical knowledge of species’ geographical and elevational distributions is therefore a high priority.

An accurate understanding of small mammal biodiversity is particularly imperative in the Madrean region given their potential to serve as zoonotic disease vectors (e.g., hantaviruses: Astorga et al. 2025; lung fungi: Salazar-Hamm et al. 2022) and indicators of ecosystem health (Pearce and Venier 2005; Leis et al. 2008; Russo et al. 2021). Understanding how many mammal species live in a certain area and how their distributions are influenced by environmental changes can help predict the risk of pathogen spillover into human populations (Olival et al. 2017; Johnson et al. 2020; García-Peña and Rubio 2024). Unfortunately, substantial gaps exist in mammal distributional knowledge across this region. This “dearth of published data” on mammal biodiversity was highlighted for the Madrean Archipelago *ca.* 20 years ago (Koprowski et al. 2005: 412), with particular sampling gaps noted for native species of bats, rodents, and shrews as compared to larger ungulates and carnivores. While the endangered Mount Graham Red Squirrel (*Tamiasciurus fremonti grahamensis*) has been intensively studied (e.g., Merrick et al. 2021; Mahoney and Pasch 2024), most small mammal species in this region are comparatively unknown. The efforts of Findley et al. (1975) and Hoffmeister (1986) in assembling the *Mammals of New Mexico* and *Mammals of Arizona*, respectively, still represent the best distributional knowledge for many Madrean Sky Island species despite their outdated taxonomies. Advances in the digitization of museum specimen records over the past two decades now present an outstanding opportunity to revisit prior work on the distribution of Madrean Sky Island small mammals, aiming to set a baseline for future studies.

We seek to address three related questions. First, what is the most functional definition of the Madrean Sky Islands for woodland-dwelling species; i.e., how many separate mountaintop islands does this system contain? Varying answers have been given to this seemingly simple question, calling for a specific geospatial definition. Second, where in the Madrean Sky Islands have small mammal species been historically studied versus not? Museum voucher specimen records are the most reliable means of addressing this question, given that small mammals are cryptic, mostly nocturnal, and therefore difficult to identify from photographs. Third, which mountains are historically understudied relative to their size? Here we present evidence of the need for targeted field surveys to holistically sample voucher specimens needed to fill geographic, elevational, and taxonomic gaps of small mammals in the Madrean Sky Islands.

## Methods

### Study system

To define the Madrean Sky Islands study system, we used NASA’s ASTER Global Digital Elevation Model (DEM) within ARCGIS Pro v3.1.2 to delimit all areas above 1,600 meters (m) in the study region. This elevational cutoff was based on Moore et al. (2013) and roughly corresponds to the lowest elevation of Madrean oak-woodland habitat. The 1,600-m cutoff served as a starting point to isolate the forested mountain areas south of the Superstitions, east of the Baboquivaris, west of the Alamo Hueco and Big Hatchets, and north of the Sierra de Las Guijas. We then performed manual cleanup to refine this delimitation, as follows. First, we eliminated mountain ranges that have less than 1 kilometer squared (km^2^) area above 1,600 m (e.g., Sierra El Humo and Sierra San Juan). Then, we determined whether contiguous areas of land above 1,600 m constituted separate ranges in the context of the dispersal propensity of small mammals. To resolve any boundary conflicts, we used satellite imagery (Google Earth; Airbus imagery 5/21/2022–11/19/2023) to identify areas of discontinuous oak woodland coverage. For example, while the Galiuro and Winchester Mountains remain continuous in terms of area above 1,600 m, a significant gap in oak woodland habitat between them constitutes separating these sky islands. Small (<1 km^2^), discontinuous fragments of area above 1,600 m were assigned to the nearest sky island if they were within 5 km of each other; otherwise these fragments were eliminated from the study. This distance is based on the estimated maximum natal dispersal distances of non-volant small mammals (Sutherland et al. 2000; Whitmee and Orme 2013). Using these criteria, we defined 54 isolated forest habitats as constituting the sky islands of the Madrean system and formalized this definition as individual DEM raster and vector polygon layers (Supplementary Data SD1-SD2; all supplementary data files are available via Github along with relevant R code: https://github.com/uphamLab/RiveraUpham_madreanSmallMammals).

Within this 54-mountain definition, we made one notable exception to separately retain the Canelo Hills and Huachuca Mountains, even though they are continuously connected in terms of both land area above 1,600 m and woodland habitat. The Huachucas reach elevations high enough to maintain pine and mixed conifer forests while the Canelo Hills comprise almost exclusively of oak woodland without elevations above 2,000 m. We reasoned that combining historical records from the steep and heterogenous Huachucas with those from the lower and more uniform Canelo Hills would be misleading. The stark difference in ruggedness as well as the dense historical sampling of the Huachucas relative to the few Canelo Hills specimens also supported this decision.

### Specimen records

We summarized natural history specimen records from public databases to assess the historical sampling efforts of small mammals in the Madrean Sky Islands. Searches of the Global Biodiversity Information Facility (GBIF) database (searched 1/23/24) allowed us to maximize the number of collections and specimen records included in the study. We focused our searches upon the taxonomic orders of Rodentia, Chiroptera, and Eulipotyphla using a geographic boundary polygon that encompassed the entire study region (bounded at four points: 33.76114, - 111.7944; 27.05179, −113.06772; 27.00817, −105.62368; 34.39024, −106.02152). This approach returned 122,324 total specimens (bats: n=21,118; rodents: n=99,844; shrews: n=1,362), including those collected in both lowland and highland habitats. We intersected all specimen records with the polygons of 54 mountains in the Madrean Sky Islands to filter records to areas above 1,600 m, which reduced the number of specimens to 10,339 total (Fig. 2; bats: n=2,260; rodents: n=7,900; shrews: n=179). We also excluded specimens with a coordinate uncertainty of 10 km or greater to maximize both the amount and accuracy of data included in this study, which further reduced the number of specimens to 9,801 (bats: n=2,194; rodents: n=7,428; shrews: n=179). This approach struck a balance between retaining known highland species while excluding known lowland species. Without this uncertainty-based filtering, the coordinates of a strictly lowland species might mistakenly be interpreted to the highest reaches of a mountain (e.g., *Chaetodipus penicillatus* at 2,500 m in the Huachuca Mountains, MSB:Mamm:184809).

**Fig 2.**
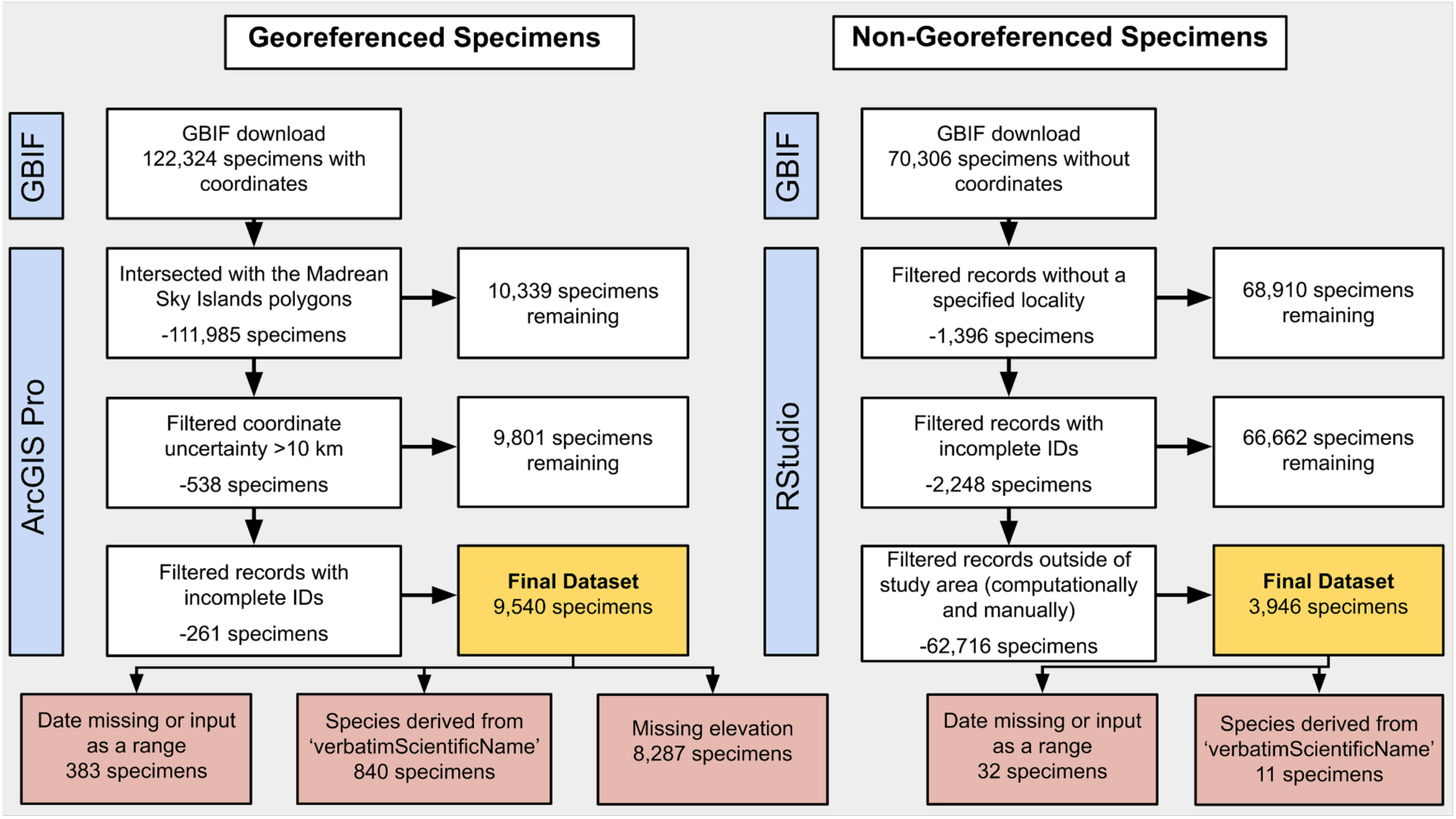
Summary of steps taken to derive the georeferenced and non-georeferenced specimen datasets (gold) used in the study along with the programs used to perform filtering (blue). Noteworthy aspects of records retained in the datasets are identified at the bottom (red), including lacking a valid date, missing elevation, or a species name derived from GBIF’s ‘verbatimScientificName’ field. Non-georeferenced specimens were identified as likely occurring in the Madrean Sky Islands by: (i) ‘computationally’ using the sky island names as search terms of the locality field and exporting only records that matched before (ii) ‘manually’ confirming the remaining records to ensure each specimen was actually captured on or near the mountains in the study area.

We used another filtering step to remove specimens that were only identified at the genus level, given that this lack of taxonomic certainty may indicate reliability issues with the record. Some specimens lacked a specific identification in GBIF’s ‘Species’ data field, but did contain a complete specific identification in the ‘verbatimScientificName’ field. For these specimens (n=840, most from University of Michigan Museum of Zoology), we added their verbatim identification to the ‘Species’ data field for inclusion in our study. Among the remaining specimens, most lacked any elevation data (n=8,287). To resolve this, we interpolated elevation using the ASTER DEM layer, performed a sensitivity test comparing interpolated and reported elevation, and favored the interpolated value even among specimens with reported elevation to ensure consistency in subsequent analyses (discrepancies between interpolated and reported elevation were not used as a criterion for removing specimens). We also tested the sensitivity of the 1,600 m cutoff by altering the elevational limit by +/- 200m in the Santa Catalinas, one of the most densely sampled sky islands. At both 1,400 and 1,800 m, we measured how the number of species, number of specimens, and area of the sky island changed relative to 1,600 m. For graphing specimen collection over time, we excluded specimens that lacked a date altogether or had been assigned a range of dates spanning multiple years (n=383; typically listed as either January 1st, 1800 or January 1st, 1900). Finally, we updated the taxonomy of the specimens to match that of the Mammal Diversity Database v2.2 (MDD 2025), which included minor spelling changes (e.g., *Erethizon dorsatus* to *Erethizon dorsatum*), re-assigned genera (e.g., *Aeorestes cinereus* to *Lasiurus cinereus*) and synonymizations (e.g., *Corynorhinus rafinesquii pallescens* to *Corynorhinus townsendii*; Supplementary Data SD3.

Further, we investigated each species to determine if existing literature generally supported the range of the species within the Madrean Sky Islands. If the known range of the species included only a part of this region (e.g., *Peromyscus truei*), the species was included in total species counts but extralimital records were flagged as in need of further investigation. If the known range of the species fell well outside of the Madrean Sky Islands, the species was not included in the total species counts and records were also flagged in the data tables.

Lastly, while recent digitization efforts have improved the resolution of specimen records online, there remain many non-georeferenced specimens (Dunnum et al. 2018). To mitigate this, we separately queried the GBIF database (searched 10/22/25–11/4/25) to identify records in Arizona, New Mexico, Sonora, and Chihuahua that lacked coordinates entirely. Subsequently, we filtered these records to the same requirements as the georeferenced specimens and searched the locality field for records matching specific sky islands. We used the name of each mountain range as search terms, with two exceptions: (i) “Graham” as an additional keyword for the Pinaleños due to its prevalence in the localities of older specimens (“Pinaleño” was rarely used to refer to this mountain range); and (ii) “Portal” as an additional keyword for the Chiricahuas due to its frequent use in the localities of specimens captured at or near the Southwestern Research Station, which lies above 1,600 m in pine-oak woodland of these mountains. The resulting specimens were manually curated to remove specimens that were still likely outside of the Madrean Sky Islands (e.g., Chiricahua Ranch on the San Carlos Indian Reservation). These specimens were not included in the main analyses due to the lack of formal georeferencing but are discussed in more detail below.

### Sampling bias

To identify potential biases of where historical collectors have focused their efforts in the Madrean Sky Islands, we evaluated the ruggedness, area, and elevational profile of each mountain. We calculated the Terrain Ruggedness Index for the individual DEMs, which returned the number of raster pixels in each of 6 ruggedness categories (level, nearly level, slightly rugged, intermediately rugged, moderately rugged, and highly rugged; Riley et al. 1999). We used these pixel counts to generate a weighted average and standard deviation of ruggedness per sky island. To quantify the specimen-area ratio (s/km^2^) as a measure of “sampling density”, we first intersected the individual polygons with the filtered rodent, bat, and shrew specimen records to identify historical specimens from each sky island individually. Then, we used the attribute table for each mountain’s DEM to calculate the area, multiplying the total number of pixels >1,600 m for each sky island by the size of one pixel in m^2^ (adjusted for longitude and latitude). Lastly, we compared the elevational profiles of mountains using the DEMs to quantify the amount of land area (km^2^) per unit elevation (m). We applied five vegetation categories to these elevational profiles, following the guidance of Bennett et al. (2004): oak woodland from 1,600-1,800 m; pine-oak woodland from 1,801-2,200 m; pine forest from 2,201-2,550 m; mixed conifer forest from 2,551-3,000 m; and limber and alpine fir forests >3,000 m (only found in the Pinaleño Mountains). We used these vegetation categories to estimate the total land area available to each habitat type across the sky islands, excluding sky islands that contained less than 1 km^2^ of a given habitat category. Also excluded from these analyses was the highly variable chaparral habitat that occurs between 1,250 m and 2,590 m depending on mountain slope and aspect.

## Results

### Study system

Using our definition of the Madrean Sky Islands as containing 54 unique mountain ranges with woodland habitats above 1,600 m, we find that this region encompasses 13,210 km^2^ of isolated habitat within a sea of desert in the Southwestern United States and Northern Mexico (Fig. 1; Table 1). The international border roughly bisects these sky islands, with 25 mountains lying primarily in the state of Sonora (SO) compared to 24 in Arizona (AZ) and 5 in New Mexico (NM). The highest elevations in the Madrean Sky Islands are found at the peaks of the Pinaleño Mountains (AZ) while the largest sky island by area is the Chiricahua Mountains (AZ). The smallest sky island by both maximum elevation and area is Cerro Bacoachi (SO).

Comparison of elevational profiles (Fig. 3) reveals that each mountain range varies greatly in slope, aspect, maximum elevation, and total area. For example, the Peloncillo South Mountains (max. elevation=2,095 m, area=921 km^2^) contain extensive oak woodland habitat and limited pine-oak woodland, but fail to reach the higher elevations needed for mixed conifer forests. On the other hand, the Rincon Mountains (max. elevation=2,646 m, area=265 km^2^) are less than one third of the area of the Peloncillo South Mountains, yet reach high enough elevations to support mixed conifer forests. In terms of habitat variation, even though the Chiricahua Mountains (max. elevation=2,990 m, area=1,752 km^2^) are 90% larger than the Peloncillo South Mountains overall, the latter contain 48 km^2^ more oak woodland.

**Fig 3.**
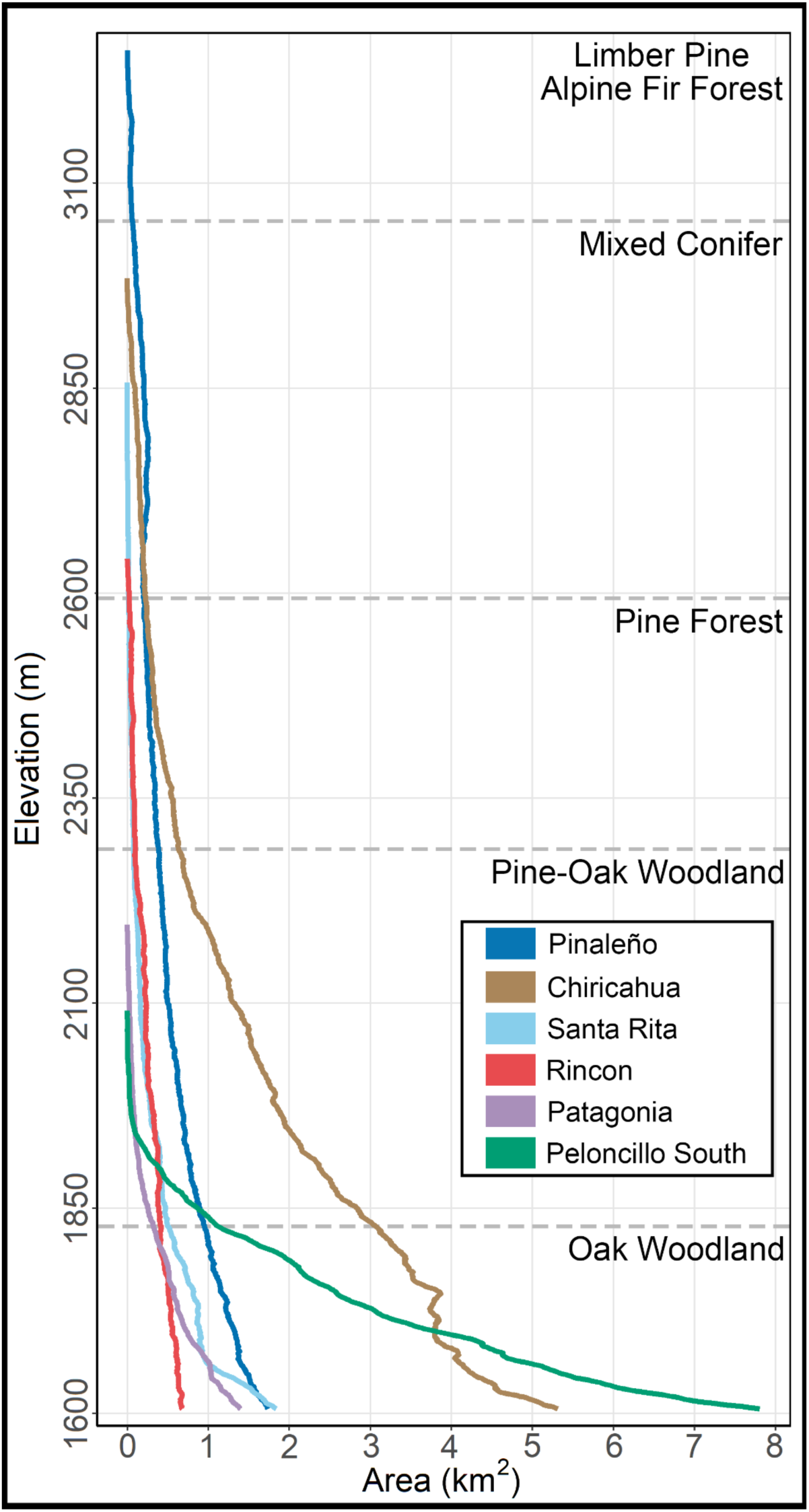
Elevation per unit land area > 1,600 m for six mountains in the Madrean Sky Island region of Arizona. Five ecoregions are overlaid with their typical upper range limit relative to elevation (sensu Bennett et al. 2004).

### Specimen records

After filtering georeferenced specimen records for quality and locational accuracy, we find that 9,540 specimens of small mammals have been collected in the Madrean Sky Islands (Fig. 4). Only 25 of the 54 mountains have one or more historical specimen records. Of those, 7,634 specimens were collected in Arizona, 1,761 in New Mexico, and 145 in Sonora (no specimens were collected in the small Chihuahuan regions of the Madrean Sky Islands). In the order Rodentia, we identify 7,227 specimens comprising 6 families (Cricetidae, Erethizontidae, Geomyidae, Heteromyidae, Muridae, and Sciuridae), 22 genera, and 50 species (Fig. 5). These specimens were collected from 25 sky islands. In the order Chiroptera, we identify 2,135 specimens comprising 5 families (Molossidae, Mormoopidae, Natalidae, Phyllostomidae, and Vespertilionidae), 20 genera, and 35 species (Fig. 6). These specimens were collected from 20 sky islands. Under the order Eulipotyphla, we identify just 178 specimens from 1 family (Soricidae), 2 genera, and 3 species (*Notiosorex crawfordi*, *Sorex arizonae*, *and Sorex monticolus*) collected from 9 sky islands (see Supplementary Data SD4-SD6).

**Fig 4.**
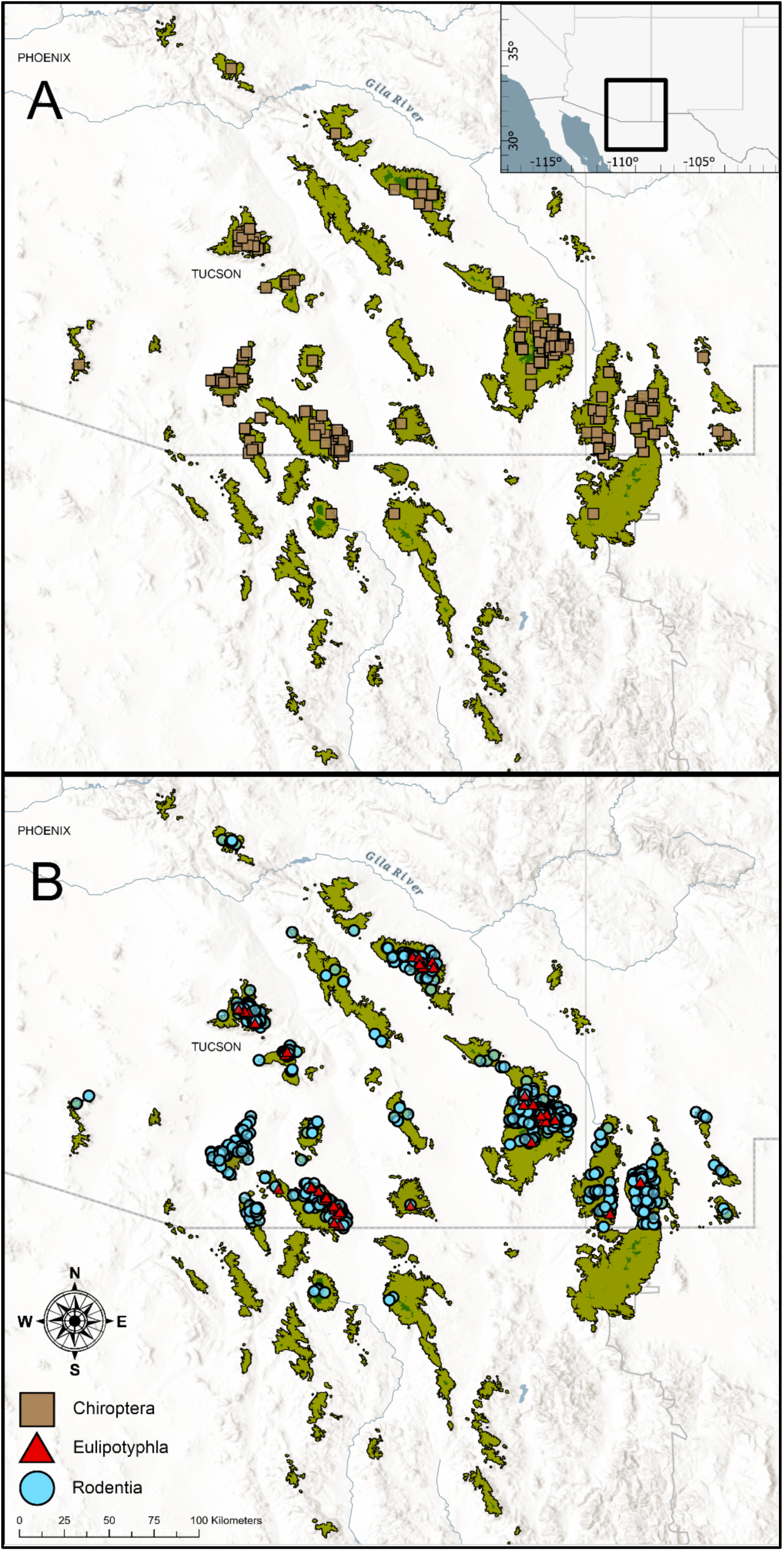
Historical voucher specimens sampled from areas > 1,600 m in the Madrean Sky Islands, as separated by (A) Chiroptera, and (B) Eulipotyphla and Rodentia.

**Fig 5.**
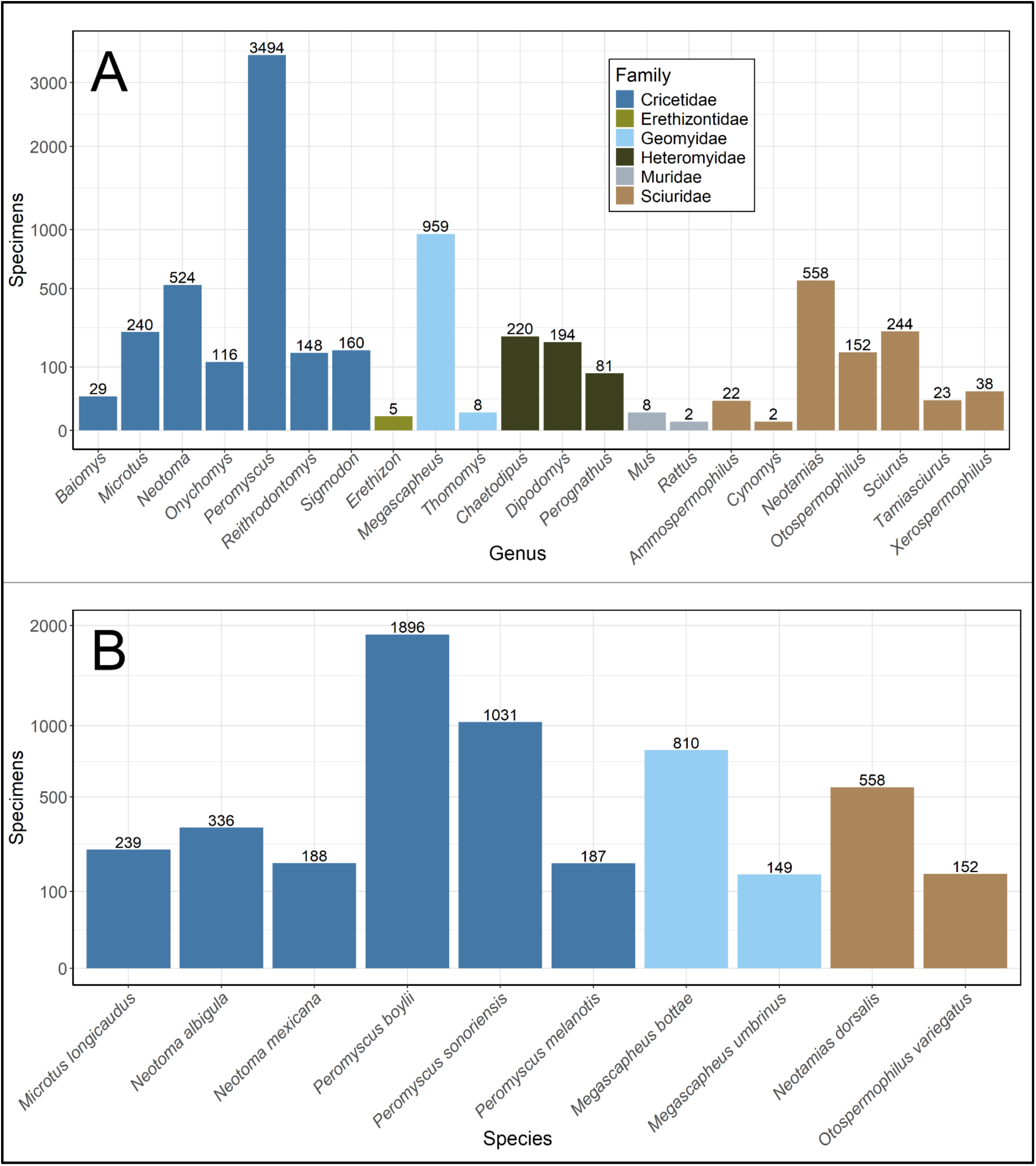
Frequency of rodent specimens sampled at > 1,600 m in the Madrean Sky Islands (A) by genus with colors corresponding to taxonomic families, and (B) by the 10 most frequently sampled species. Both charts are on a square root scale.

**Fig 6.**
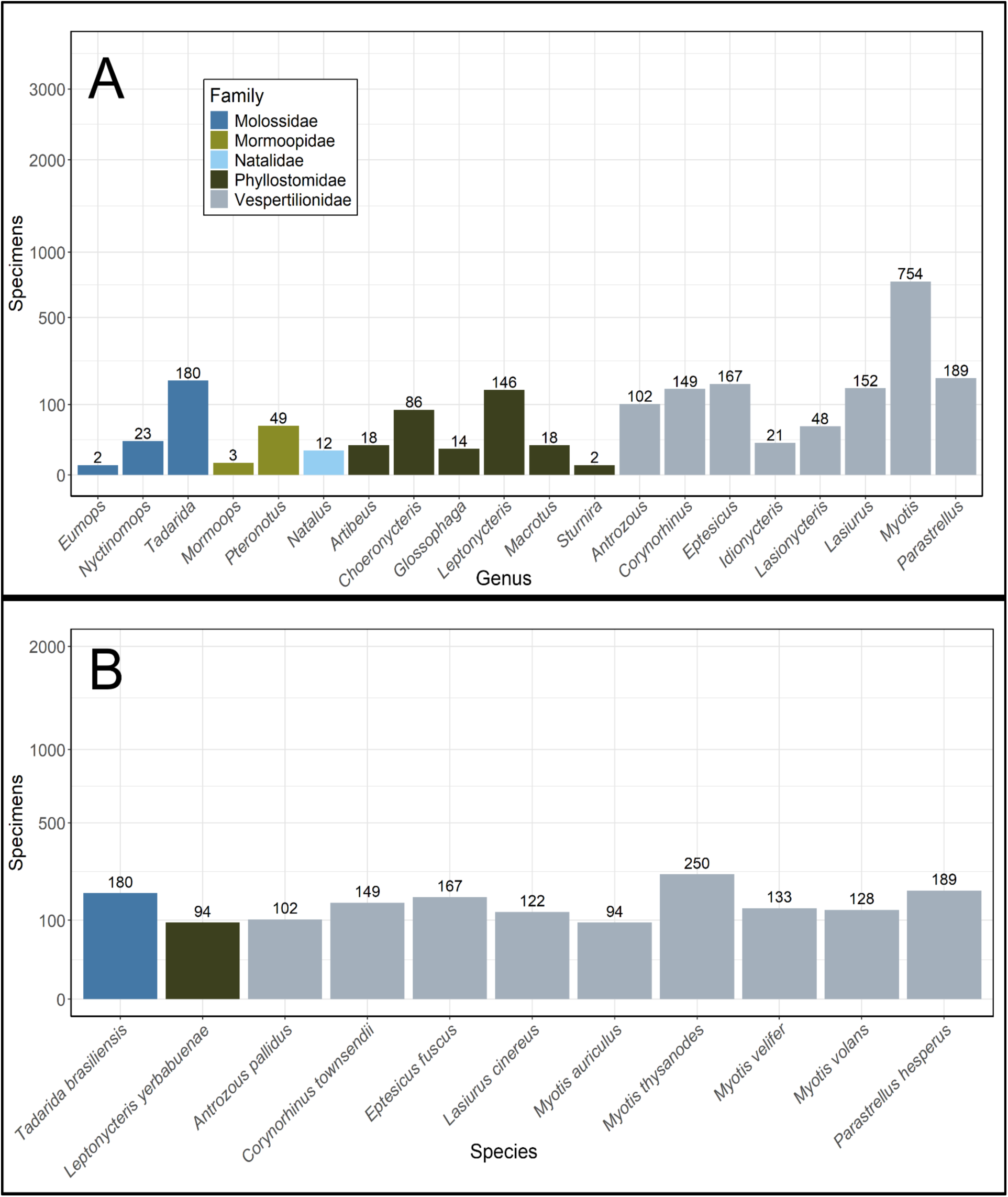
Frequency of bat specimens sampled at > 1,600 m in the Madrean Sky Islands (A) by genus with colors corresponding to taxonomic families, and (B) by the 10 most frequently sampled species (*Leptonycteris yerbabuenae* and *Myotis auriculus* are equally the 10th most common). Both charts are on a square root scale.

Considering the timing of collecting activity in the Madrean Sky Islands, the earliest specimen was collected in the Huachuca mountains by E. A. Mearns in 1884 (FMNH:Mammals:4943) and only 265 specimens were collected across the sky islands before 1900. The majority of specimens (5,921; 65%) were collected between 1950 and 1980, encompassing 84% of bats, 59% of rodents, and 48% of shrews ever collected in the Madrean Sky Islands (Fig. 7; Supplementary Data SD7). In the ensuing 43-year period (1981-2023), only 1,142 specimens were collected in this region (12% of the total).

**Fig 7.**
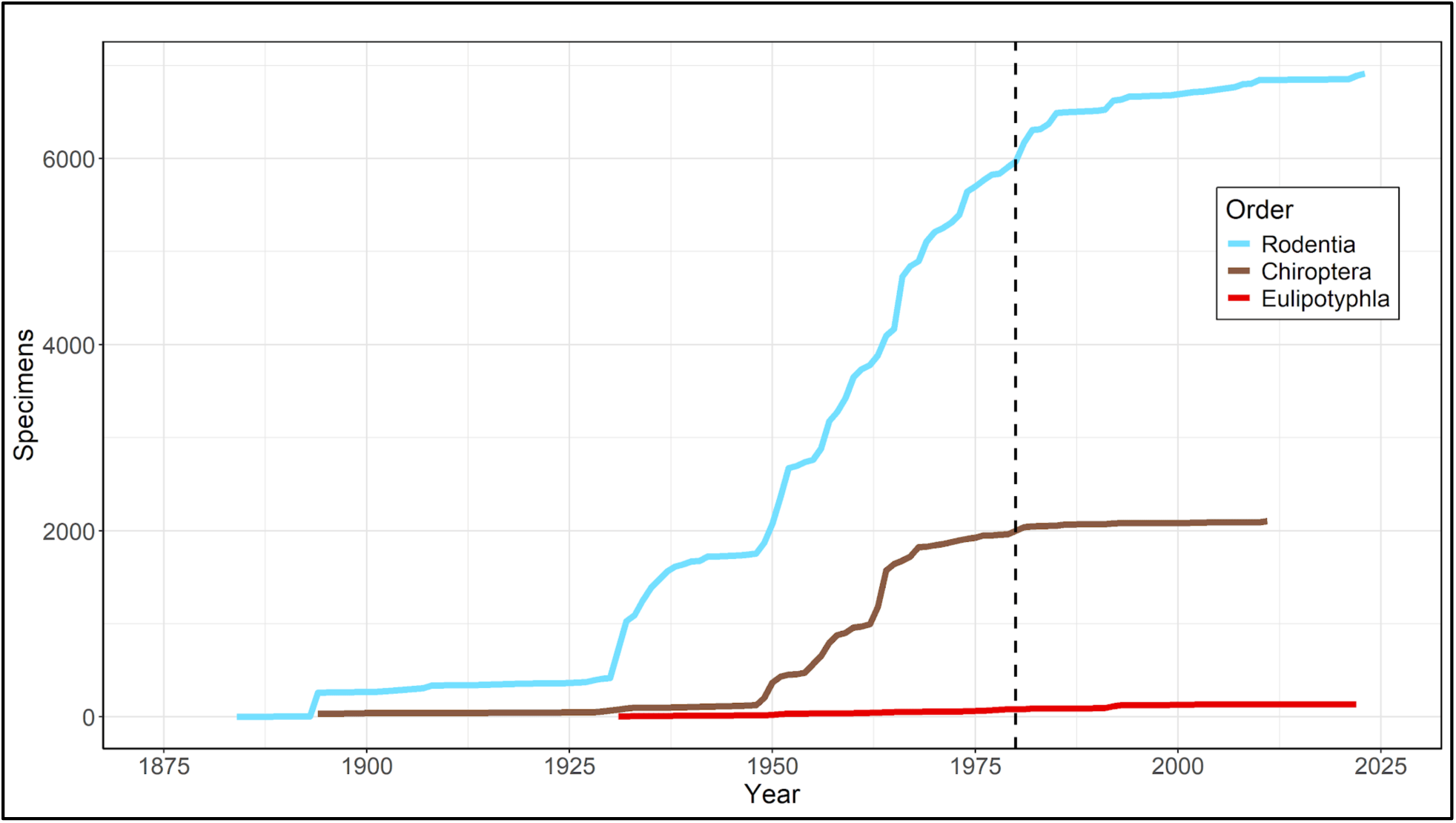
The accumulation of specimens over time across Chiroptera, Rodentia, and Eulipotyphla. The colored lines begin where the first specimen for each order was collected (1884, 1894, and 1931 respectively) and the vertical dashed line indicates the year 1980.

**Fig 8.**
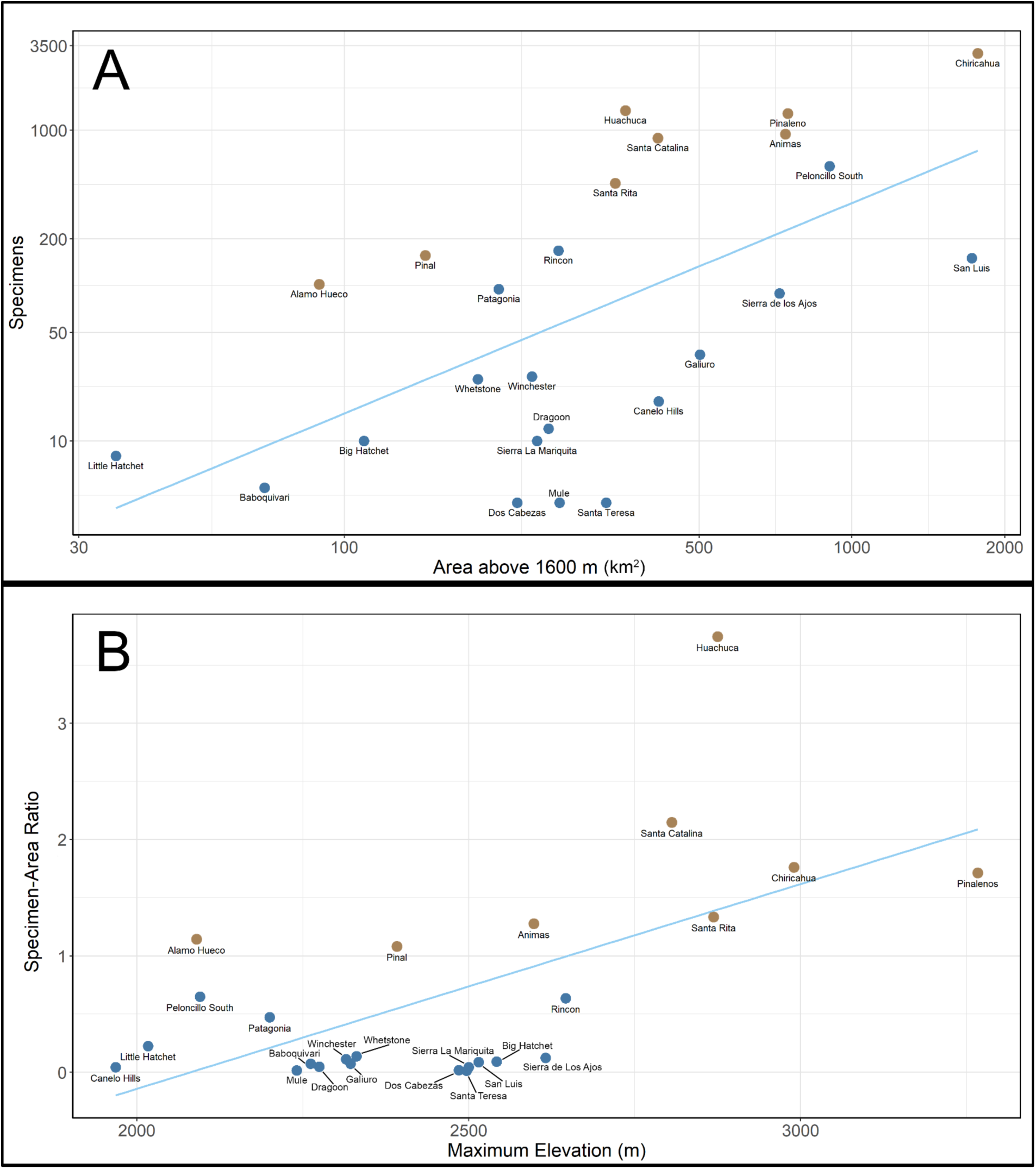
(A) Biases in historical sampling as measured by the number of small mammal specimens relative to mountain area > 1,600 meters. Both axes are shown on a log10 scale, indicating that larger sky islands have disproportionately more specimens than smaller ones. (B) Biases in historical sampling as measured by the density of sampling relative to sky island maximum elevation on a linear scale. In both panels, the 29 sky islands without any georeferenced specimens were excluded and sky islands with a specimen-area ratio greater than 1 are shown in brown.

Preserved specimens of small mammals from the Madrean Sky Islands are presently housed in 43 different natural history collections, mainly in the southwestern United States (increases to 64 collections if non-georeferenced specimens are included, see next paragraph). Only 3 georeferenced specimens from the study area are housed in Mexican museums, with 1 from the Instituto de Biología de la UNAM (IBUNAM) and 2 from the Centro de Investigaciones Biológicas del Noroeste (CIBNOR). The most extensive collections are the Museum of Southwestern Biology (University of New Mexico) with 3,377 specimens, followed by the University of Arizona Museum of Natural History (1,954 specimens), University of Michigan Museum of Zoology (1,090 specimens), and Arizona State University Natural History Collection (394 specimens).

Our analysis of non-georeferenced specimens yielded an additional 3,964 records lacking coordinates that are deemed to likely fall within the boundaries of our study area. Of these, 122 specimens were collected in Mexico and 3,842 were collected in the United States. Six additional species were recorded: *Chaetodipus goldmani*, *Dipodomys deserti*, *Megascapheus fulvus*, *Perognathus amplus*, *Eumops underwoodi*, and *Nyctinomops femorosaccus*. A total of 24 sky islands were associated with non-georeferenced specimens, 6 of which lacked georeferenced specimens in our prior analyses: Pajaritos, Sierritas, Sierra Azul, Sierra de Oposura, Sierra San Jose, and Sierra San Antonio. The earliest non-georeferenced specimen was collected in 1855 and only 316 specimens were collected after 1980 (representing 8% of the non-georeferenced specimens). Non-georeferenced specimens are held in 42 different collections (21 of which are unique from those that hold georeferenced specimens), including the Smithsonian National Museum of Natural History (1,342 specimens), the American Museum of Natural History (786 specimens), the Gila Center for Natural History at Western New Mexico University (532 specimens), and the Museum of Southwestern Biology (311 specimens). IBUNAM is the only Mexican museum with non-georeferenced specimens, represented by 4 records (see Supplementary Data SD8).

### Sampling bias

We find that sky island area and maximum elevation significantly influence where historical collectors focused their sampling. Collection efforts leading to the preservation of small mammal specimens have been concentrated in a few, large sky islands in the Madrean system, resulting in a positive log-log relationship between sky island size and specimens (Fig. 7A; log(*Y*) = −3.524 + 1.354 log(*X*), *R*^2^=0.3578, *P*=0.0016). Similarly, we find a positive linear relationship between the maximum elevation of a sky island and the specimen-area ratio (Fig. 7B; *Y* = −3.664 + 0.0017 *X*, *R*^2^=0.3762, *P*=0.0011). Interestingly, we found no significant relationship between the specimen-area ratio and sky island ruggedness (*R*^2^=0.0791, *P*=0.1731), even though ruggedness is positively correlated with maximum elevation (*R*^2^=0.2901, *P*=0.0055). The average specimen-area ratio across all 54 sky islands is 0.32 s/km^2^, or 0.68 s/km^2^ after excluding the 29 mountains without specimens (Supplementary Data SD9-SD10). The most densely sampled sky islands are the Huachucas (3.74 s/km^2^), Santa Catalinas (2.15 s/km^2^), Chiricahuas (1.76 s/km^2^), Pinaleños (1.71 s/km^2^), and Santa Ritas (1.33 s/km^2^); only 3 other sky islands have >1.0 s/km^2^ (the Animas, Alamo Hueco, and Pinal Mountains). We recover notably low sampling density on several large sky islands: the San Luis Mountains (2,515 km^2^, 0.09 s/km^2^), the Sierra de los Ajos (735 km^2^, 0.12 s/km^2^), Galiuros (503 km^2^, 0.07 s/km^2^), and Canelo Hills (423 km^2^, 0.04 s/km^2^).

The addition of non-georeferenced specimens to each mountain does not significantly alter these results. The relationship between sky island area and number of specimens when both georeferenced and non-georeferenced specimens are included is represented by the line log(*Y*) = −1.210 + 0.995 log(*X*) (*R^2^*=0.271, *P*=0.003). Overlapping 95% confidence intervals indicate this slope is not significantly different than when georeferenced-only data are used (slope interval of 0.692–1.299 versus 0.976–1.732 using the georeferenced-only data). The relationship between the maximum elevation of a sky island and its specimen-area ratio including both sets of specimens is represented by the line *Y* = −3.778 + 0.002 *X* (*R^2^*=0.247, *P*=0.004). Once again, overlapping 95% confidence intervals indicate that including non-georeferenced specimens does not significantly change the results (0.0014–0.0026 versus 0.0013–0.0023 using the georeferenced-only data).

We tested the accuracy of our DEM-interpolated elevations for the 1,253 georeferenced specimens that also have a reported elevation. We find the relationship between reported and interpolated elevation to have a slope of 0.451 (*R^2^*=0.503, *P*<0.001). However, this relationship seems to be strongly influenced by specimens which have the same value in both the ‘elevation’ and ‘elevationAccuracy’ fields, possibly indicative of errors in data entry. By excluding these 123 specimens from the analysis, the slope changes to 0.778 (*R^2^*=0.705, *P*<0.001; Supplementary Data S11). An additional 15 specimens have >1,000 m discordance between interpolated and reported elevation, presumably for reasons other than data entry error; we nevertheless decided to use the interpolated value for consistency across all specimens. We also tested the effects of altering the 1,600 m cutoff by +/- 200 m in the Santa Catalina Mountains due to its high sampling density and isolation from other sky islands. At 1,600 m, the Santa Catalinas have 890 small mammal specimens across 42 species, a total area of 414.78 km^2^, and a sampling density of 2.15 s/km^2^. We find that reducing the cutoff to 1,400 m increases the number of specimens to 999, adds three species (*Chaetodipus intermedius*, *C. baileyi*, and *Dipodomys spectabilis*), increases the total area to 511 km^2^, and decreases the sampling density to 1.96 s/km^2^. Increasing the cutoff to 1,800 m decreases the number of specimens to 864, removes one species relative to 1,600 m (*Perognathus flavus*), decreases the total area to 270.07 km^2^, and increases the sampling density to 3.20 s/km^2^ (Supplementary Data S12).

## Discussion

The dearth of published research highlighted by Koprowski et al. (2005) has remained a reality for most species of small mammals in the Madrean Sky Islands. However, thanks to 2 decades of advances in the digitization of museum specimen records, we are now able to synthesize data from 64 natural history collections to identify gaps in the distributional knowledge of this diverse small mammal fauna. The ‘virtual flora and fauna’ of Deyo et al. (2013) was the last attempt to summarize distributional knowledge of taxa across the Madrean Sky Islands, but the surprisingly few records of mammals (n=217) they recovered was likely an artifact of sparse digitization and georeferencing efforts at that time. Their work preceded the maturation of integrated databases like VertNet and GBIF (Guralnick et al. 2016; Heberling et al. 2021), which now enable greater synthesis of historical specimen records. In contrast, our search and quality-filtering steps revealed 9,540 georeferenced voucher specimens of small mammals (Fig. 3). These specimens are confidently allocated to 88 species from 12 families of bats, rodents, and shrews (Figs. 4, 5). The over 40x growth in digitized records from this region illuminates several biases in historical collecting, including at least 23 of the 54 mountains in the Madrean Sky Islands for which no small mammals have ever been vouchered (29 unsampled mountains considering only georeferenced vouchers).

To put this sampling history in regional context, a study by Jones et al. (2021) in the neighboring Gila Mountains region of New Mexico found 12,505 specimen records from 77 species of bats (n≈2,500), rodents (n≈9,250; totals estimated from figures), and shrews across nearly twice the study area (24,383 km^2^ versus 13,210 km^2^ in the Madrean Sky Islands). Both regions experienced similar peaks of collecting activity prior to 1980, but the large pulse of Gila collecting from 2012–2020 (led by the Museum of Southwestern Biology) differs from the comparatively dormant collecting in the Madrean Sky Islands over the same period (Fig. 6 compared to fig. 6c in Jones et al. 2021). Critically, the paucity of post-1980 specimens from the Madrean Sky Islands means that frozen tissues and holistic specimen preparations are mostly unavailable (both preservation modes were popularized subsequently; Galbreath et al. 2019, Timm et al. 2025). Overall, Madrean small mammals have been less extensively studied than those of the Gila Mountains despite the former’s ∼2x higher density of species per km^2^ (0.0067 versus 0.0032). Recent efforts to re-survey the Santa Catalina Mountains using holistic preservation (2021–2023 led by the Arizona State University Natural History Collections; see Rowsey et al. this volume) have begun bridging this shortfall, but parallel efforts on nearby sky islands are sorely needed.

Digitally defining the mountains of the Madrean Sky Islands was a key step in this study that now enables comparable future studies in this biodiverse region (Fig. 1). Querying how woodland-dwelling taxa of different kinds have responded to recent climatic and land use changes in this region is now more replicable given the included geospatial resource (Supplementary Data SD1-SD2). However, the 54-mountain definition of the Madrean Sky Islands presented here differs from three previous totals: “about 40” in Warshall (1995: 6); “55 ranges/complexes” in Deyo et al. (2013: 296); and “∼65 mountain ranges” in Moore et al. (2013: 148). Those differences, and some nuances of our own definition, underscore the inherent subjectivity of dividing a complex, rugged landscape into discrete units. The extent of the forest in this region is also fluctuating, and has changed even over the ∼150-year period of specimen collecting summarized in this study (Yanahan and Moore 2019). We followed the lead of Moore et al. (2013) in choosing an initial definition of 1,600 m, corroborating that definition with a finding that lower elevational thresholds introduced more desert-adapted, heteromyid rodents and included habitat not relevant to most woodland-dwelling species. Then, we deviated from that work by joining several of the mountains that they acknowledged were part of ‘Madrean Sky Island complexes.’ The result better reflects the ability of woodland-dwelling small mammals to disperse through continuous oak woodland (Sutherland et al. 2000; Whitmee and Orme 2013).

For example, we joined the Patagonia and San Antonio Mountains, which in Moore et al. (2013) appear to only be separated by the U.S.-Mexico international border (Supplementary Data SD13). In the case of the mountain complex consisting of Chiricahua, Pedregosa, Swisshelm, and Dos Cabezas, these ranges are connected by varying degrees of oak woodland and so under that criterion could be considered one unit (as Warshall 1995 chose to do). We chose to join the Chiricahua and Pedregosa Mountains owing to their elevational and forest connectivity above 1,600 m while separating the Swisshelm and Dos Cabezas Mountains since they lack such elevational connectivity. Satellite imagery does show sparse oak woodland growth below 1,600 m connecting these latter two mountains from the Chiricahuas (Google Earth; Airbus imagery 5/21/2022–11/19/2023), but we reasoned that this thin forest likely presents a strong enough dispersal barrier to warrant separate mountain recognition. While this definition of the Madrean Sky Islands includes some grassland areas adjacent to oak woodland (e.g., indicated by specimens of several heteromyid species), it remains a useful approximation of the time-averaged area that species in oak woodlands and above have been inhabiting during the study period.

We found clear patterns of uneven among-mountain sampling across the Madrean Sky Islands that makes accurate small mammal biodiversity assessments difficult. Past collecting efforts have concentrated mainly on 8 sky islands that collectively contain 87% of known rodent, bat, and shrew specimens. These more densely sampled mountains all contain >1.0 specimens/km^2^ (Fig. 7A), consistent with the expectation of positive specimen-area relationships if larger mountains attract more collecting efforts. However, the comparatively sparse sampling of other large sky islands (e.g., San Luis and Sierra de los Ajos in Mexico, Galiuros and Canelo Hills in the U.S.) indicates that mountain area is not the only factor explaining historical collecting. Indeed, the maximum elevation of a mountain is also strongly associated with increased sampling density (Fig. 7B), perhaps in part owing to the development of paved roads to areas above 1,600 m on some of the taller mountains (e.g. the Pinaleños, Santa Catalinas, and Santa Ritas). Road accessibility, a known bias in biodiversity surveys (Kadmon et al. 2004; Monsarrat et al. 2019), was likely critical in promoting surveys of mountains with established roads. Yet roads may also threaten biodiversity by promoting land conversion (Bennett 2017; Barrientos et al. 2021). Thus, sky island access roads may paradoxically be associated with both increased biodiversity knowledge (as measured in specimens and resulting publications) and potential threats, a tradeoff that land managers must closely consider.

Even for sky islands with comparatively dense historical sampling, small mammal diversity is still likely underestimated. For example, the Huachuca Mountains have the highest sampling density of any Madrean Sky Island at 3.67 s/km^2^, yet over a century of fieldwork on the mountain failed to find the Cliff Chipmunk (*Neotamias dorsalis*) until a breeding population was identified in 2007 (Cudworth and Koprowski 2010). Furthermore, a 2023 field survey of the Santa Catalina Mountains (2.15 s/km^2^) identified the first specimen of the Fulvous Harvest Mouse (*Reithrodontomys fulvescens*) on that sky island (Rowsey et al. this volume). We suggest that the status of the Madrean Sky Islands as a ‘biogeographic crossroads’ (Spector 2002) raises the likelihood that additional species at the northern or southern edge of their distributions have gone undetected on multiple mountains. Intensively surveying elevational gradients across adjacent sky islands may present a useful test of the hypothesis that species have ‘rare edges and abundant cores’ (Brown 1984; Martin et al. 2024). Future work that mitigates the identified sampling biases for small mammals may be able to accomplish this goal.

Comparing elevational profiles among sky islands shows striking heterogeneity in the extent of different forest types across mountains (Fig. 2), suggesting that biodiversity knowledge gained from one sky island might not apply to all others. For example, the Peloncillo South and Patagonia Mountains have similar maximum elevations (2,095 and 2,200 m, respectively) but the former contains ∼4.5x more forested area than the latter across a sloping oaken plateau. These disparate elevational profiles are associated with more total specimens (587 vs. 95, respectively) and more apparent species diversity in the former than the latter (27 rodent, 19 bat, and 1 shrew species compared to 12, 5, and 0). While this positive correlation between sky island area and species richness is consistent with Frey et al. (2007), no systematic method exists for identifying which species will be lost and which will be retained as the sky island area shrinks. Hence, we expect each additional sky island for which the resident small mammal fauna is surveyed adds value to the total biodiversity knowledge of the Madrean system.

One of the main limitations to our study is the unevenness in the extent of specimen digitization and georeferencing among natural history collections. This could be partially responsible for the sparse sampling of Mexico’s Madrean Sky Islands, particularly the collections from smaller, regional museums that may still be digitally inaccessible (Dunnum et. al. 2018). However, 2 of the largest museums in Mexico have extensive mammal collections published on GBIF and still lack Madrean Sky Islands representation: (i) CIBNOR has digitized 13,025 small mammal specimens (all georeferenced), only 2 of which are in the sky islands from 1,673 in the initial download polygon; and (ii) IBUNAM has 48,688 small mammal specimens (34,650 georeferenced) and only 1 in the sky islands of 642 in the initial download. A more pervasive issue across the Madrean Sky Islands appears to be specimens that are digitized, but that lack coordinate data. Notably, our search of georeferenced specimens on GBIF failed to return records from the American Museum of Natural History (AMNH) and the Smithsonian National Museum of Natural History (NMNH), two large historical collections for which georeferencing efforts are incomplete. By querying GBIF specifically for records that had not been georeferenced, we found an additional 3,964 specimens that were likely captured within the bounds of our study area, including 1,342 and 786 specimens respectively held at the NMNH and AMNH. Most were collected on the same mountains for which georeferenced data were available, such that their inclusion in our dataset did not change any main findings. We highlight that, although extensive georeferencing is beyond the scope of our study, such efforts to map historical specimen localities will be extremely valuable for enabling fine-grained studies (Bloom et al. 2018).

Considering specimens with geospatial coordinates (georeferenced or recorded by GPS) in the Madrean Sky Islands, the accuracy of digital records and the historical methods used to collect them present two further challenges. First, while online biodiversity data aggregators like GBIF have advanced the capacity to synthesize specimen records globally (Heberling et al. 2021), GBIF data requires careful validation to correct errors and outdated taxonomies (Zizka et al. 2020). Because a majority of the specimens in our study’s initial download predate the availability of handheld GPS devices (Kumar and Moore 2002), their latitude and longitude presumably came from secondary georeferencing using field notes and locality descriptors. Our filtering based on coordinate uncertainty thresholds aimed to alleviate unreliable records, but this approach still relies on georeferencing accuracy (Murphey et al. 2004). Discordance between interpolated and reported elevation for some specimens (Supplementary Data S11) highlights this limitation and suggests that the coordinate and elevation were occasionally derived independently during the georeferencing effort. Relying on either elevation value for further analyses without ground-truthing the actual habitat of that site requires some caution, although we reasoned that the interpolated values are obtained consistently across all specimens and thus more reliable overall. Second, the lack of transparency regarding which trapping techniques were used to capture specimens on each sky island presents another bias. For example, mountains surveyed using Museum Special snap traps may yield greater diversity and abundance of small mammals per trapnight than those using Sherman live traps (Eulinger and Burt 2011). The unevenness of bat-netting efforts is another potential bias; however, bat specimens were recovered from most of the same mountains as non-volant taxa (Fig. 3) and bats and rodents have roughly parallel specimen accumulation curves through time (Fig. 6). The apparent rarity of shrews in museum collections could reflect the infrequent use of pitfall traps, drift fences, and other methods (Maddock 1992), rather than actual low abundances in the Madrean Sky Islands.

Taxonomic changes and historical difficulties in identifying small mammals present other challenges to this study. The external measurements that are often used to differentiate related species can be prone to error, potentially leading to misidentification (Blackwell et al. 2006). Indeed, in our review of specimen records, we identified 11 species whose identification requires further investigation (Supplementary Data SD14). For example, the 18 specimens identified as *Leptonycteris curasoae* in the Madrean Sky Islands are presumably either *L. nivalis* or *L. yerbabuenae*, given that *L. curasoae* is endemic to Colombia and Venezuela (Cole and Wilson 2006), but making this determination would require physically examining those specimens (all housed in the MSB Mammal Collection). Similarly, our dataset includes 1 specimen of *Peromyscus beatae*, a species of deer mouse native to central Mexico with no known distribution in the Mexican portion of the Madrean Sky Islands (Álvarez-Castañeda et al. 2017), indicating that this specimen may be a misidentified individual of *P. boylii* (Tiemann-Boege et al. 2000). So too with the 11 specimens of *P. truei* that are recorded from sky islands in Arizona (Chiricahuas and Santa Catalinas) when their known distribution is thought to be restricted to the Animas Mountains of New Mexico south into the Sierra Madre Occidental (Hoffmeister 1981; Cook 1986). Investigation is needed to determine if these cases constitute modern range expansions, historical undersampling, or previous misidentifications. These issues highlight the steadfast importance of museum specimens as physical records, each one providing a critical opportunity to revisit observations, collect new data (e.g., measurements, DNA sequences), and thereby gain new insights even hundreds of years later.

Overall, we find that woodland-dwelling small mammals are surprisingly understudied throughout the Madrean Sky Islands. These forested mountains exist at the crossroads of the Nearctic and Neotropical biogeographic realms, which presents a diverse setting of repeated forest isolation and reconnection for studying how ecological and evolutionary processes unfold (Marshall 1957, Warshall 1995, Spector 2002, Koprowski et al. 2005, Deyo et al. 2013, Moore et al. 2013). We find that only 8 of the 54 mountains have had substantial historical sampling of voucher specimens, and at least 23 mountains are still unsampled. The multiplicity of threats now impacting this region from fires, drought, and human disturbance signals an imperative to document basic biodiversity patterns across the sky islands (Misztal and Hansen 2013; Peters et al. 2018; Yanahan and Moore 2019; Love et al. 2023). Geopolitical conflicts such as the construction of a steel wall along the US-Mexico international border, bisecting as many as six sky islands, also impact biodiversity in this region. This wall directly blocks dispersal routes for larger mammals and indirectly impacts smaller mammals via increased light pollution, human presence, and road infrastructure (Harrity et al. 2024; Marín and Koprowski 2025). Increased fire severity, notably in more mesic and high-elevation biomes, has also played a role in the degradation of sky island habitat, which has the potential to decrease diversity among small mammal communities (O’Connor et al. 2014; Villarreal et al. 2019; Culhane et al. 2022).

Therefore, future studies should focus on elevational sampling in un- and under-sampled sky islands, particularly in Mexico where only two mountains (San Luis, Sierra de Los Ajos) have georeferenced records of >10 small mammal specimens. So too should future studies focus on cryopreserving tissues and preparing holistic specimens (*sensu* Galbreath et al. 2019, Rowsey et al. this volume) to better investigate host-pathogen interactions and the microbiome more broadly, two fields in which wild species are under-represented (Pascoe et al. 2017, Wu et al. 2018). Even the 8 comparatively well-sampled sky islands mostly lack post-1980 collection efforts necessary for such studies, corroborating a broader trend in the decline of mammal specimen collection since the mid-20^th^ century (Patterson 2002, Malaney and Cook 2018). Land managers and state agencies should take the evidence of biodiversity knowledge shortfalls in the Madrean Sky Islands as a call to establish baseline collections where they are lacking and promote re-survey efforts when possible, enabling studies of how and why small mammal communities are changing through time.

### Conservation and Management Implications

Though the results of this study indicate that the whole of the Madrean Sky Islands region is substantially undersampled, several sky islands emerge as high priorities to sample and preserve holistic voucher specimens in the near future. In Arizona, the Galiuro Mountains and Canelo Hills are the largest and most undersampled ranges and should thus be highest priority. The Galiuros are surrounded by the Santa Catalinas and Pinaleños and the Canelo Hills are adjacent to the Huachuas, making both targets ideal for contextualizing existing specimens from nearby, densely-sampled mountains. In New Mexico, the Big Hatchet Mountains have the lowest density of historical specimens, so sampling them would provide key insights on small mammal communities in smaller mountains of the Madrean Sky Islands. Conversely, the San Luis Mountains form the second largest sky island of this system and cross multiple borders in Sonora, Chihuahua, and New Mexico, representing a politically challenging but fascinating mountain to study small mammal diversity. The scant sampling of the San Luis make them a high priority and a key opportunity for developing cross-border biodiversity collaborations.

A future in which multiple, thoroughly sampled elevational gradients in the Madrean Sky Islands are established will enable replicated tests of which factors are the most important determinants of species richness (Chen et al. 2017) and metacommunity structure (Presley et al. 2012), two long-standing eco-evolutionary questions. Extending these comparisons to small mammal viromes, myco-, and micro-biomes will additionally be possible only if high-quality preservation of voucher specimens and flash-frozen tissues are prioritized, opening new doors for studying phylosymbiosis and codiversification (e.g., Suzuki et al. 2022, Chen et al. 2023), among other questions. Finally, comparing the population-genetic histories of forest-dwelling small mammals in this system will enable studies of how differing dispersal abilities affect gene flow between conspecific populations on different mountains. Apt candidates for this research include some of the most common sky island species in museum collections, such as *Peromyscus boylii* and *Corynorhinus townsendii*.

## Data availability

All data and code used to generate these analyses are posted on Github here: https://github.com/uphamLab/RiveraUpham_madreanSmallMammals/

## Acknowledgments

We owe thanks to the generations of mammalogists that have contributed invaluable specimens from which we derive small mammal biodiversity knowledge in the Madrean Sky Islands. Dakota M. Rowsey helped to inspire and guide this work. Special thanks to Luisa Zamora Chavez, Alexander Fenlon, Gilma De Leon, Jessica Neumeier, Ryan Nguyen, Morgan Pierce, Julia Nitschmann and two anonymous reviewers for helpful comments that improved this manuscript. Funding support was provided by the National Institutes of Health (1R35GM156919 to NSU) and Arizona State University, as well as the American Society of Mammalogists (Grant-in-Aid of Research to DCR).

## Supplementary Data

Supplementary Data SD1.—54 individual sky island vector polygon shape layers and supporting files for use in GIS software.

Supplementary Data SD2.—54 individual sky island DEM raster layers and supporting files for use in GIS software.

Supplementary Data SD3.—List of taxonomic changes made to the original GBIF names.

Spelling changes and synonymizations were made to match the Mammal Diversity Database v2.2. Taxonomic changes that resulted in range shifts were resolved only when the name could be unambiguously assigned to one other species, otherwise, they were left as is and discussed in Supplementary Data S14. For example, *Glossophaga soricina* once included *G. mutica* along with several other species but is now restricted to South America and *G. mutica* is the only species found in North America. Specimens only identified to the genus level by GBIF but containing a species ID in the ‘verbatimScientificName’ field are also included here.

Supplementary Data SD4.—Table of Chiroptera specimens included in the study with original GBIF data fields as well as columns added during the study (e.g., Species_MDD).

Supplementary Data SD5.—Table of Eulipotyphla specimens included in the study with original GBIF data fields as well as columns added during the study (e.g., Species_MDD).

Supplementary Data SD6.—Table of Rodentia specimens included in the study with original GBIF data fields as well as columns added during the study (e.g., Species_MDD).

Supplementary Data SD7.—Combined table of all specimens included in the study with the exception of specimens that lacked a date altogether or had been assigned a range of dates spanning multiple years (n=383).

Supplementary Data SD8.—Table of non-georeferenced specimens included in the study with original GBIF data fields as well as columns added during the study (e.g., sky_island)

Supplementary Data SD9.—Data matrix summarizing area, mean ruggedness, maximum elevation, and specimen counts for each sky island.

Supplementary Data SD10.—Raw area and ruggedness data for each sky island.

Supplementary Data SD11.—(A) Comparison between reported and interpolated elevation values for the 1,253 specimens that have a reported elevation. Elevation was interpolated by overlaying the specimen coordinates over a digital elevation model. The black line represents *Y=X* and the blue line represents the slope of the linear regression. (B) Comparison between reported and interpolated elevation values after removing specimens (n=123) that had the same value in both the ‘elevation’ and ‘elevationAccuracy’ fields in GBIF, possibly indicating an error during data entry. The slope of the relationship between the two values increases from 0.451 to 0.778.

Supplementary Data SD12.—Specimens included in the Santa Catalina Mountains at three different thresholds (1,400, 1,600, and 1,800 m).

Supplementary Data SD13.—Changes made to the mountains included in the Madrean Sky Islands compared to Moore et al. 2013.

Supplementary Data SD14.—List of 11 species with specimen identifications found in the Madrean Sky Islands that need further examination. These observations fall outside of the currently understood ranges of these species. Both *P. truei* and *P. difficilis* are known to occur on specific sky islands, however, some specimens have been reported outside of these known mountains, warranting additional verification. Specimens of *L. ega* and *M. keenii* likely belong to *L. xanthinus* and *M. evotis* respectively, but verification of specimens is also needed.

